# Targeting mechanistic target of rapamycin complex 2 attenuates immunopathology in Systemic Lupus Erythematosus

**DOI:** 10.1101/2024.08.01.606069

**Authors:** Minji Ai, Xian Zhou, Michele Carrer, Paymaan Jafar-nejad, Yanfeng Li, Naomi Gades D.V.M., Mariam Alexander, Mario A. Bautista, Ali A. Duarte Garcia, Hu Zeng

## Abstract

**Objective:** We aim to explore the role of mechanistic target of rapamycin complex (mTORC) 2 in systemic lupus erythematosus (SLE) development, the in *vivo* regulation of mTORC2 by type I interferon (IFN) signaling in autoimmunity, and to use mTORC2 targeting therapy to ameliorate lupus-like symptoms in an *in vivo* lupus mouse model and an *in vitro* coculture model using human PBMCs.

**Method:** We first induced lupus-like disease in T cell specific *Rictor*, a key component of mTORC2, deficient mice by topical application of imiquimod (IMQ) and monitored disease development. Next, we investigated the changes of mTORC2 signaling and immunological phenotypes in type I IFNAR deficient Lpr mice. We then tested the beneficial effects of anti-*Rictor* antisense oligonucleotide (*Rictor*-ASO) in a mouse model of lupus: MRL/*lpr* mice. Finally, we examined the beneficial effects of *RICTOR*-ASO on SLE patients’ PBMCs using an *in vitro* T-B cell coculture assay.

**Results:** T cell specific *Rictor* deficient mice have reduced age-associated B cells, plasma cells and germinal center B cells, and less autoantibody production than WT mice following IMQ treatment. IFNAR1 deficient Lpr mice have reduced mTORC2 activity in CD4^+^ T cells accompanied by restored CD4^+^ T cell glucose metabolism, partially recovered T cell trafficking, and reduced systemic inflammation. *In vivo Rictor*-ASO treatment improves renal function and pathology in MRL/*lpr* mice, along with improved immunopathology. In human SLE (N = 5) PBMCs derived T-B coculture assay, *RICTOR-*ASO significantly reduce immunoglobulin and autoantibodies production (P < 0.05).

**Conclusion:** Targeting mTORC2 could be a promising therapeutic for SLE.

## Introduction

Systemic lupus erythematosus (SLE) is an autoimmune disease characterized by loss of tolerance to self-antigen and the production of autoantibodies. An estimated of 0.40 million people are diagnosed with SLE annually among whom over 85% are young female patients globally (1). SLE is known for its heterogeneity. It presents various clinical manifestations, from mild skin rashes to life-threatening multi-organ damages such as nephritis, cerebritis, and myelofibrosis (2). Broad immunosuppressive agents such as azathioprine are commonly used for SLE management which may not lead to optimal disease control and can lead to undesired side effects (3). Unlike rheumatoid arthritis with numerous disease-specific biologics, only two biological agents, belimumab and anifrolumab, are approved for SLE treatment (4). Therefore, a better understanding of SLE pathogenesis is urgently needed for the development of novel and specific therapeutic agents.

SLE is a heterogeneous disease influenced by various factors including genetic, environmental, and immunological factors (5). Multiple molecular pathways, particularly type I interferon (IFN) signaling, have been proposed to play pivotal roles in disease pathogenesis (6). Over-activation of plasmacytoid dendritic cells (pDCs) leads to elevated IFNα and induced genes, the type I IFN signature, in SLE patients (7), which promote the presentation of self-antigens to autoreactive T and B cells leading to autoimmunity (8). Injection of IFN-αβ to NZB/W F1 mice has led to the rapid onset of lupus-like diseases (9). Conversely, IFN-I receptor-deficient mice were partially protected from pristane induced lupus mice (10), lupus-prone NZB mice (11) and C57BL/6-Fas*^lpr^*mice (12). Although one report indicated that deficiency of type II but not type I IFN receptor ameliorates lupus-like diseases in MRL/*lpr* mice (13), treatment with anti-IFNAR blocking antibody alleviated lupus-like symptoms in MRL/*lpr* mice (14), suggesting a disease promoting function of type I IFN in MRL/*lpr* model. Monoclonal antibodies directed at the type I IFN receptor, such as Anifrolumab, have been actively tested in clinical trials for SLE treatment and showed promising outcomes (15). Despite the well-known link between type I IFN signature and SLE, there are several outstanding questions in this context, including the mechanisms through which type I IFN signaling modulates T and B cell functions in lupus. We found previously that activation of type I IFN can synergize with TCR signaling to promote the mechanistic target of rapamycin (mTORC) 2 in T cells *in vitro*, which may contribute to Tfh cell-mediated immunopathology, and T cell lymphopenia in lupus (16). Whether such IFN/ mTORC2 axis mediates lupus *in vivo* remains unknown.

The mechanistic target of rapamycin (mTOR) is an evolutionally conserved serine/threonine kinase complex regulating various cellular processes including growth, survival, and metabolism. mTOR has two forms of complexes, mTORC1 and mTORC2, which play indispensable but distinct roles in T cell biology. mTORC1 is critical for naïve T cell activation, proliferation, and effector T cell differentiation (17), while mTORC2 mediates Tfh cell differentiation (18). In regulatory T cells, mTORC1 maintains natural or thymic-derived Treg while overactivation of mTORC2 suppresses Treg function and the ability of Tfh cell inhibition (19). Thus, mTORC2 activities regulate Tfh and Treg balance which can be critical in autoimmune diseases such as SLE. We previously showed that the genetic deletion of mTORC2 in CD4^+^ T cells in C57BL/6-Fas*^lpr^* (Lpr) mice, a lupus-prone mice, showed improved immunopathology associated with reduced Tfh differentiation and glucose metabolism (16). Here, we provide evidence that mTORC2 deficiency can ameliorate TLR7 agonist induced lupus development, which is associated with high type I IFN signature and dependent on pDC (20). We also found that type I IFN receptor deficient Lpr mice have reduced mTORC2 activities linking to ameliorated lupus-like symptoms. Finally, we showed that pharmacological targeting mTORC2 can benefit lupus-like mice *in vivo* and reduce autoantibodies production in an *in vitro* co-culture system using T and B cells from SLE patients. Together, our results support that targeting mTORC2 in T cells could be a therapeutic option for SLE.

## Material and Method

### Mice

*Cd4*^Cre^*Rictor*^fl/fl^ mice have been described before (18). *Ifnar1*^−/−^ (Strain #: 028288), C57BL/6-Fas*^lpr^* (Lpr) (Strain #: 000482) and MRL/MpJ-Fas*^lpr^*/J (MRL/*lpr*) (Strain #: 000485) mice were purchased from the Jackson Laboratory. *Ifnar1*^−/−^ mice were crossed with C57BL/6-Fas*^lpr^* mice to generate C57BL/6-Fas*^lpr^Ifnar1*^−/−^ mice. A total of 82 mice were used in this study, with a mix of male and female mice on *Cd4*^Cre^*Rictor*^fl/fl^ background, and female only mice on Fas*^lpr^* background. MRL/*lpr* mice used for therapeutic testing were randomly allocated into different experimental groups. Mice were housed under specific pathogen-free conditions on a 12:12-h day: night cycle with access to normal chow (LabDiet, 5P76) and water. The room temperature was 22 ± 1 °C, with 31% humidity. All mice were bred and maintained in the Department of Comparative Medicine at Mayo Clinic Rochester. All animal procedures were approved by the Institutional Animal Use and Care Committee (IACUC).

### Human sample collection/storage

Five SLE patients (53.6 ± 4.15 years) and five age-matched healthy donors’ (55 ± 5.01 years) blood samples were collected for peripheral blood mononuclear cells (PBMC) isolation. PBMCs were isolated using Ficoll density gradient centrifugation and cryopreserved in liquid nitrogen until use. SLE patients fulfill the EULAR/ACR SLE 2019 criteria (21). Patients’ demographics, disease manifestations and medications were detailed in Supplementary Table 1. The disease activity index was assessed using the Systemic Lupus Erythematosus Disease Activity Index 2000 (SLEDAI-2K) based on the manifestations presented one month before sample collection (22).

### ASO Treatment

Ionis Pharmaceutical manufactured anti-mouse *Rictor*-ASO and anti-human *RICTOR*-ASO and corresponding Ctrl-ASO. Each mouse received *Rictor*- or Ctrl-ASO subcutaneously at 50 mg/kg once per week. Mice were treated for 6 consecutive weeks. Mouse proteinuria was measured weekly by Urine Reagent Strips (Siemens).

### Flow Cytometry

Single cell suspension of spleen and peripheral lymph nodes was prepared as previously described (18). Cells were first stained with Fixable Dye Ghost 510 (Tonbo Bioscience) for viability, then with desired surface marker antibodies on ice for 30 mins. Antibodies used are detailed in supplementary methods. FACS data was acquired by the Attune NxT (ThermoFisher) cytometer. Data analysis was performed in FlowJo software (Tree Star).

### ELISA

ELISA was used to detect serum autoantibody levels. Briefly, 96-well plates (2596; Costar) were coated with dsDNA (2 µg/ml in PBS), ssDNA (2 µg/ml in PBS), or chromatin (Sigma-Aldrich, 5 µg/ml) overnight at 4°C. Plates were washed 4 times with PBS-T (0.05% Tween 20 in PBS) and blocked with 5% blocking protein (Bio-Rad) at 37°C for 1 h. Plates were then washed 4 times with PBS-T before adding serially diluted serum. Serum coated plates were incubated at 37°C for 1.5h. Plates were next washed 8 times before horseradish peroxidase (HRP)-conjugated detection Abs for IgG (Bethyl Laboratories). Coated plates were incubated at 37°C for another 1.5h, washed 8 times, and added tetramethylbenzidine (TMB) substrate. The reaction is stopped by 2N H_2_SO_4_ and read at 450nm by a plate reader. Mouse serum and detection antibodies were diluted in dilution buffer (1% BSA in PBS-T).

### Immunoblotting

Mouse CD4^+^ T cells were enriched from lymph node single-cell solution using EasySep™ Mouse CD4^+^ T Cell Isolation Kit (STEMCELL, Cat # 19852). Human CD4^+^ T cells were enriched from PBMCs using EasySep™ Human CD4^+^ T Cell Isolation Kit (STEMCELL, Cat # 17952). The same numbers of cells from each sample were used for the experiment. Cells were lysed in RIPA lysis buffer (Sigma). Lysed protein concentrate was used for electrophoresis and membrane transfer. The transferred membrane was first blocked with 5% milk in TBST (0.1% Tween 20) for 1h at room temperature, washed, and incubated with primary antibodies overnight. The following primary antibodies have been used: anti-phospho-AKT (Ser473) (D9E), AKT (pan) (40D4), anti-p-S6 (Ser235/Ser236, D57.2.2E), anti-RICTOR (53A2) and anti-b-actin (13E5). The next day, the membrane was washed and incubated with corresponding secondary antibodies for subsequent enhanced chemiluminescence (ECL; Thermo Fisher) exposure. Images were captured on an Azure Imaging system. Blot intensity was quantified using ImageJ software.

### Metabolic assay

Mouse CD4^+^ T cells were enriched using EasySep™ Mouse CD4^+^ T Cell Isolation Kit (STEMCELL, Cat # 19852). Isolated cells were activated with plate-coated anti-CD3 (2 µg/ml, Bio X Cell) and anti-CD28 (2 µg/ml, Bio X Cell) for 48h, and rested overnight without stimulation. Live cells were purified with lymphocyte isolation buffer (MP Biomedicals) and then restimulated with plate-coated anti-CD3 (2 µg/ml) and anti-ICOS (5 µg/ml, Biolegend, Cat# 313502) for 24 h. Metabolic activities of ICOS-stimulated cells were then measured by the SCENITH method (23); a detailed description is provided in supplementary methods.

### T-B cell coculture assay

T-B coculture assay was performed as previously described (24). Detailed description of the assay is provided in the supplementary methods. Coculture supernatant was collected for immunoglobulin isotypes and autoantigen detection. Autoantigen microarray was performed by Genecopoeia Inc.

### Immunoglobulin isotypes detection

Mouse (IgG1, IgG2a, IgG2b, IgG3, IgA, IgM) and human (IgG1, IgG2, IgG3, IgG4, IgA, IgM) immunoglobulin isotypes were measured by LEGENDplex mouse immunoglobulin isotyping panel (Biolegend; cat# 740493) and human Immunoglobulin Isotyping Panel (6-plex) (Biolegend; cat# 740639) respectively following the manufacturer’s instructions.

### Inflammatory cytokines detection

Mouse serum cytokine level was measured by LEGENDplex Mouse Inflammation Panel (Biolegend, cat# 740446) according to the manufacturer’s instructions.

### Statistical analysis

All data are presented as mean ± SEM. One-way analysis of variance (ANOVA) with post-hoc Tukey test was used for multiple groups comparison. Unpaired and paired student t-tests was used for two groups comparison. The Kaplan-Meier survival analysis was used to compare the survival probability differences among groups. Detailed statistical tests are described in individual figure legends. Statistical analysis and graph generation were performed in GraphPad Prism version 9.0.

## Results

### Mice with T cell specific *Rictor* deletion develop less systemic inflammation upon TLR7 agonist challenge

We previously demonstrated that loss of mTORC2 in T cells substantially alleviates autoimmunity in C57BL/6-Fas*^lpr^*(Lpr) mice (16). However, Lpr mice do not develop overt kidney pathology. Toll-like receptor (TLR) 7 signaling is known to contribute to lupus initiation and exacerbation in humans (25) and mice (26). Topical application of a TLR7 agonist, imiquimod (IMQ), induces lupus-like symptoms with tissue pathology in C57BL/6 (WT) mice (20). To study whether the loss of mTORC2 signaling in T cells may impact TLR7 induced SLE development in mice, we topically administered IMQ on *Cd4*^Cre^*Rictor*^fl/fl^ mice. Following 6 weeks of IMQ application, *Cd4*^Cre^*Rictor*^fl/fl^ mice had a lower spleen weight than their WT counterparts (Figure 1A). Flow cytometry analysis of splenocytes showed that *Cd4*^Cre^*Rictor*^fl/fl^ mice had a significantly lower percentage of T-bet^+^B220^+^ and CD11c^+^B220^+^ age-associated B cells (ABCs) (Figure 1B, supplementary Figure 1A), germinal center (GC) B cells (Figure 1C) and plasma cells (Figure 1D), indicating overall attenuated B cell activation. Correspondingly, a higher percentage of naïve B cells was seen in *Cd4*^Cre^*Rictor*^fl/fl^ mice (supplementary Figure 1B). While no change of marginal zone B cells was observed (supplementary Figure 1C), a lower percentage of Bcl6 expressing B cells (supplementary Figure 1D) was seen in *Cd4*^Cre^*Rictor*^fl/fl^ mice. Additionally, there was a lower percentage of Tfh cells (Figure 1E) and CD44^hi^CD62^low^ effector CD4^+^ T cells (Figure 1F) in *Cd4*^Cre^*Rictor*^fl/fl^ mice. Little change of Foxp3^+^ Treg cells (supplementary Figure 1E) was seen between two groups of mice, while *Cd4*^Cre^*Rictor*^fl/fl^ mice had fewer T-bet expressing CD4^+^ T cells (supplementary Figure 1F). We also observed reduced plasmacytoid dendritic cells (pDCs) (supplementary Figure 1G) in *Cd4*^Cre^*Rictor*^fl/fl^ mice, a population has been reported critically linked to lupus development (27). Other myeloid cell populations including cDC, monocytes and neutrophils remained unchanged (supplementary Figure 1 G-J). These data were consistent with previous observations that mTORC2 critically contributes to humoral immunity in Lpr mice (16). They further indicate that mTORC2 in T cells could be required for the formation of ABC, a B cell lineage critical for TLR7 mutation mediated lupus pathogenesis (28). To assess systemic inflammation, we quantified inflammatory cytokines in the serum of IMQ treated mice. *Cd4*^Cre^*Rictor*^fl/fl^ mice had lower levels of cytokines such as IL1α, IFNβ, IFNγ, IL23, and IL27 (Figure 1G), commonly associated with lupus pathogenesis in both murine models and humans (29). ELISA measurements also showed that *Cd4*^Cre^*Rictor*^fl/fl^ mice had lower levels of anti-dsDNA and anti-histone antibodies than their WT counterparts (Figure 1H). Finally, kidney histology showed that WT mice developed moderate glomerulosclerosis with mesangial thickening, which were significantly improved in *Cd4*^Cre^*Rictor*^fl/fl^ mice, indicating a less extent of kidney damage (Figure 1I). Taken together, these results indicate that mTORC2 deficiency in T cells effectively impedes the immunopathologic transition in IMQ-induced lupus model.

**Figure 1:**
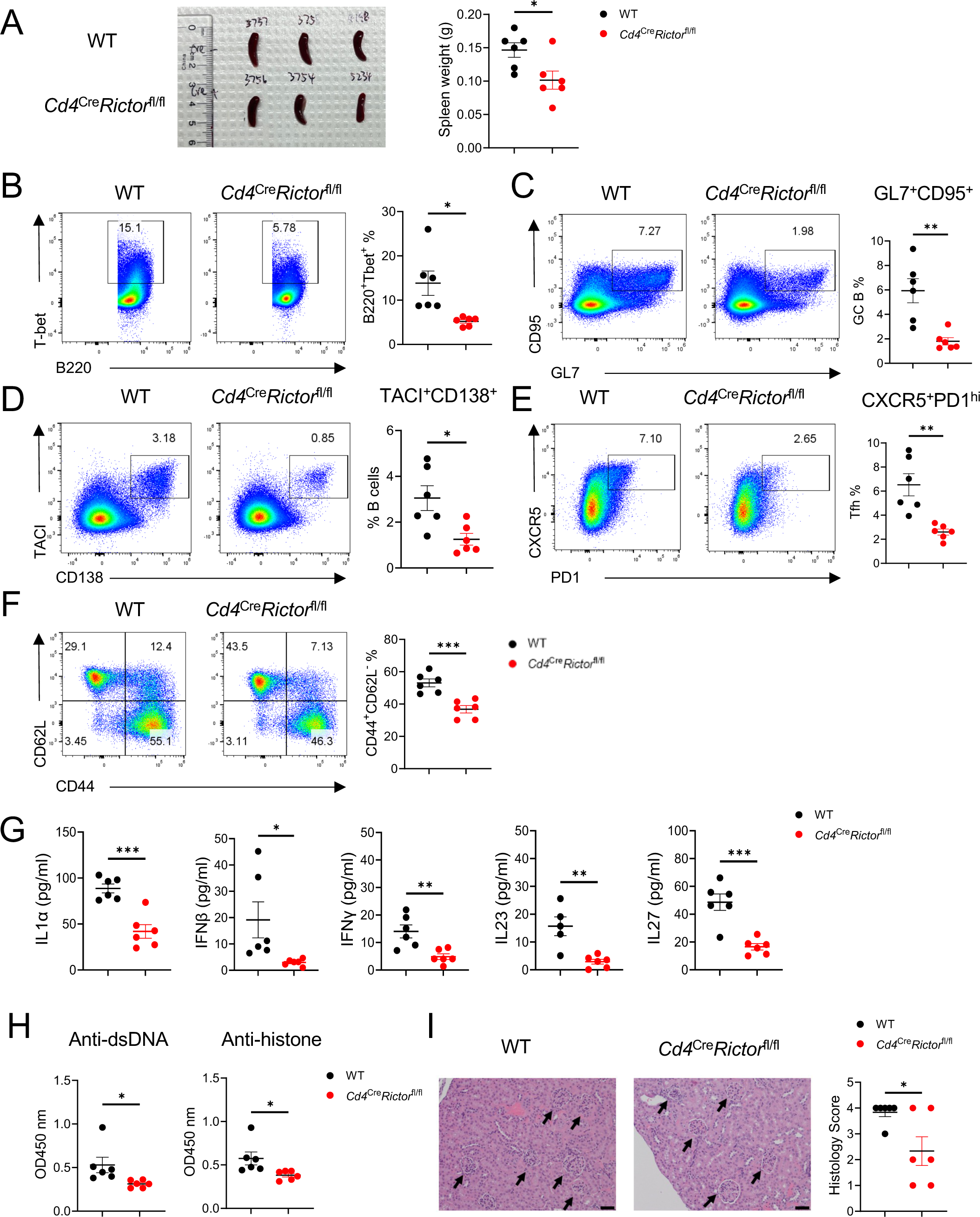
Mice with RICTOR deletion in T cells develop less severe lupus-like diseases following imiquimod (IMQ) treatment. WT and *Cd4*^cre^*Rictor*^fl/fl^ mice were treated with IMQ epicutaneously 3 times a week for 6 weeks. (A) Spleen size (left) and weight (right) of IMQ-treated mice. (B) Expression of T-bet and B220 in splenocytes. Right, summary of T-bet^+^B220^+^ age-associated B cell frequencies. (C) Expression of CD95 and GL7 on splenocytes. Right, summary of CD95^+^GL7^+^ germinal center B cell frequencies. (D) Expression of TACI and CD138 on splenocytes. Right, summary of TACI^+^CD138^+^ plasma cell frequencies. (E) Expression of CXCR5 and PD-1 on splenocytes. Right, summary of CXCR5^+^PD-1^hi^ Tfh cell frequencies. (F) Expression of CD62L and CD44 on CD4+ T cells from splenocytes. Right, summary of CD44^+^CD62L^–^ effector T cell frequencies. (G) The inflammatory cytokine levels in IMQ treated mouse serum. (H) Antibody tiers of Anti-dsDNA and anti-histone antibodies in IMQ treated mouse serum measured by ELISA. The serum was diluted at 1:100. (I) Representative H&E staining images of kidneys. Right, summary of histology scores. Scale bar: 50 µm. *p < 0.05, **p < 0.01, ***p<0.001. p-Values were calculated with unpaired t-tests. Error bars represent SEM.

### IFNAR1 deletion partially inhibits mTORC2 activities in T cells and restores TCR/ICOS-mediated glucose metabolism in Lpr CD4^+^ T cells

We previously showed that type I IFNs synergize with TCR signaling to active mTORC2 *in vitro* (16). Thus, we hypothesized that elevated type I IFN signaling in SLE contributes to mTORC2 activation *in vivo*, which promotes disease development. To test this hypothesis, we generated IFNAR1 deficient Lpr (Lpr-*Ifnar1^−/−^*) mice and investigated mTORC2 activities in these mice. To directly evaluate the mTORC2 activation in CD4^+^ T cells, we performed immunoblot on purified CD4^+^ T cells derived from different groups of mice. Elevated phosphorylated AKT (p-AKT) S473 expression, the direct target of mTORC2, was seen in Lpr T cells, which was reduced in T cells from Lpr-*Ifnar1^−/−^* mice (Figure 2A), indicating reduced mTORC2 activation in CD4^+^ T cells of Lpr-*Ifnar1^−/−^* mice. mTORC2 is known to modulate CD69 surface expression on CD4^+^ T cells (18). We previously showed that mTORC2 is required for the increased CD69 expression on Lpr T cells, which partially contributes to T cell lymphopenia phenotype in Lpr mice (16). As expected, IFNAR1 deficiency led to reduced CD69 expression in Lpr CD4^+^ T cells in blood (Figure 2B) and peripheral lymph nodes (pLN) (supplementary Figure 2A), as well as increased CD4^+^ T cell frequency in both blood (Figure 2C) and pLN (supplementary Figure 2B) in Lpr mice. These results are consistent with our hypothesis that type I IFN contributes to mTORC2 activation in CD4 T cells, and with the clinical observation that type I IFN signature is strongly associated with CD4 T cell lymphopenia in SLE patients (30). mTORC2 is also critical for TCR/ICOS-mediated glucose metabolism (18). Following TCR/ICOS stimulation *in vitro*, Lpr CD4^+^ T cells have significantly increased glycolytic capacities than WT cells, and such increase was significantly reduced in the absence of IFNAR1 (Figure 2D). Interestingly, we also observed partially restored proliferation in Lpr-*Ifnar1^−/−^* CD4^+^ T cells (Figure 2E). Because mTORC2 deletion does not restore CD4^+^ T cell proliferation in Lpr mice, type I IFN likely modulates CD4^+^ T cell proliferation independent of mTORC2 signaling. Despite the partial rescue on glucose metabolism and proliferation, IFNAR1 deletion has little rescue effect on the apoptosis of CD4^+^ T cells in Lpr mice (Figure 2F). In summary, IFNAR1 deficiency led to reduced mTORC2 activity and mTORC2-mediated T cell lymphopenia and glucose metabolism in Lpr mice.

**Figure 2:**
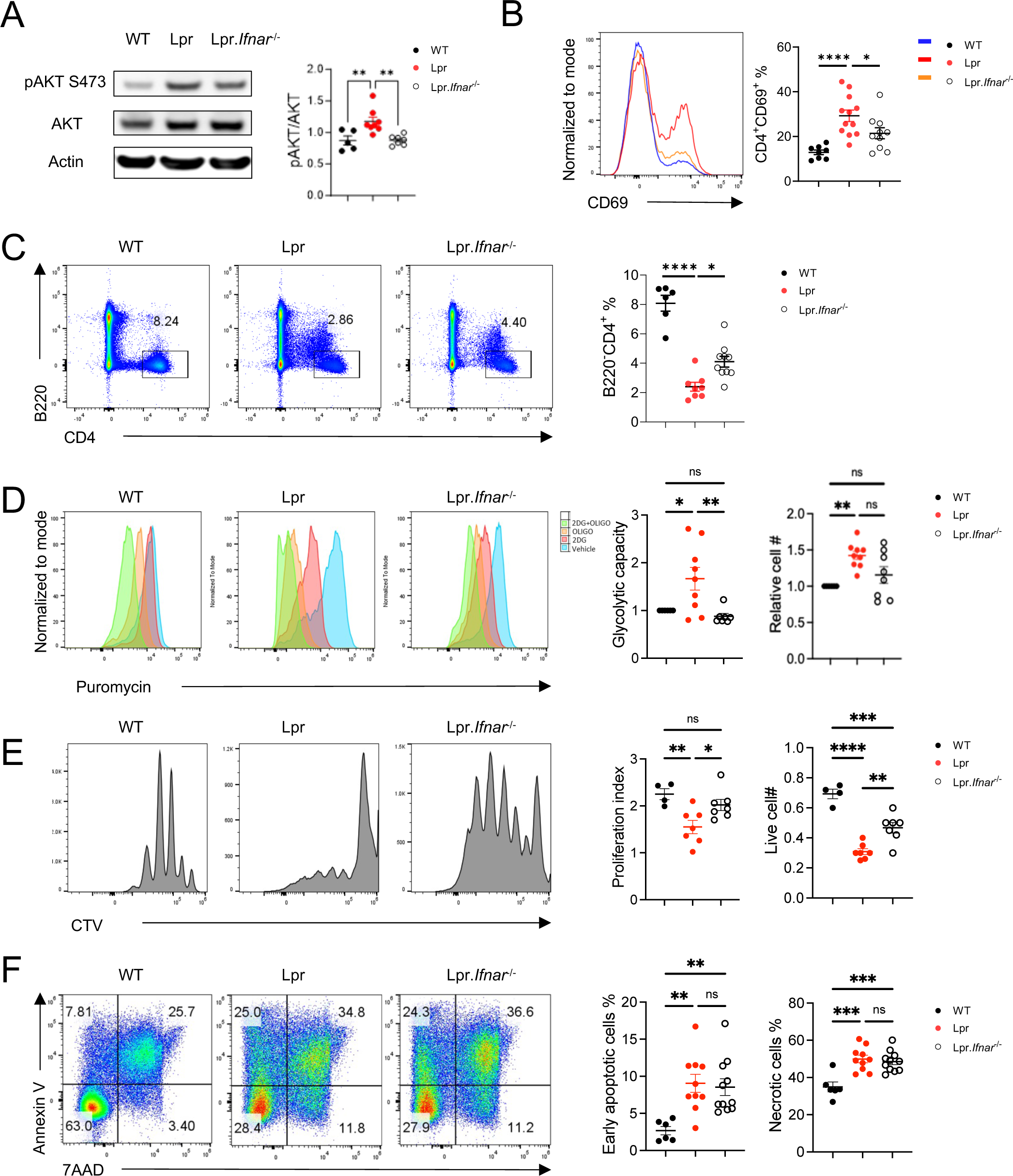
IFNAR1 deletion inhibits mTORC2 activity and rescues TCR/ICOS mediated glucose metabolism in CD4^+^ T cells of Lpr mice. (A) Immunoblot image (left) of p-AKT473 in CD4^+^ T cells derived from the lymph nodes of 6-month-old B6, Lpr and Lpr*.Ifnar1*^–/–^ *mice*. Right, relative pAKT473 expression level normalized to AKT expression in each mouse. (B) Expression of CD69 and CD4 on cells derived from peripheral blood. Right, summary of CD4^+^CD69^+^ cell frequencies. (C) Expression of B220 and CD4 in peripheral blood. Right, summary of B220^+^CD4^+^ cell frequencies. (D) Puromycin incorporation assay in CD4^+^ T cells after sequential anti-CD3/anti-CD28, and anti-CD3/anti-ICOS stimulation. Left: glycolytic capacity of CD4^+^ T cells after sequential stimulation; right: relative cell number change after sequential stimulation. Data were normalized to B6 in each experiment. (E) Representative flow plots of Cell trace violet (CTV) dilution in CD4^+^ T cells after 3 days of anti-CD3/anti-CD28 stimulation. Left: proliferation index; right: absolute cell number after stimulation. (F) Flow staining of Annexin V and 7AAD on CD4^+^ T cells after overnight stimulation of anti-CD3/anti-CD28. Left: frequency of early apoptotic cells; right: frequency of necrotic cells. 2-DG, 2-deoxyglucose; Oligo, Oligomycin. ns, not significant; *p < 0.05, **p < 0.01, ***p < 0.001, ****p < 0.0001. p-Values were calculated with one-way ANOVA with the post-hoc Tukey test. Error bars represent SEM.

### IFNAR1 deletion rescues immunopathology in C57BL/6-Fas*^lpr^* mice

Elevated interferon signature has been identified in both SLE patients (7) and lupus-prone mice (31). However, type I IFNAR deletion or inhibition yields controversial outcomes in Lpr and MRL/*lpr* mice (12,13). We revisited this question and examined the mTORC2-associated immunophenotype changes in Lpr-*Ifnar1^−/−^* mice. Compared to Lpr mice, Lpr-*Ifnar1^-/-^* mice had smaller sizes of peripheral lymph nodes and yielded a lower number of lymphocytes (Figure 3A). Lpr-*Ifnar1^−/−^* mice had lower levels of anti-dsDNA (Figure 3B) and lower concentrations of immunoglobulin isotypes including IgG1, IgG2a, and IgA than those from Lpr mice (Figure 3C). Furthermore, IFNAR1 deficiency reduced serum concentrations of multiple inflammatory cytokines, including TNFα, MCP1, IFNβ, IL17A, and IL23 (Figure 3D). The abnormal immune phenotypes presented in Lpr mice such as accumulated aberrant B220^+^TCRβ^+^ cells (Figure 3E) and CD4^-^CD8^-^DN cells (Figure 3F), modestly but significantly reduced in Lpr-*Ifnar1^−/−^* mice. Reflecting reduced autoantibodies and immunoglobulin levels, Lpr-*Ifnar1^−/−^*mice have reduced CD138^+^B220^int^ population (supplementary Figure 2C) and reduced Bcl6 expression in B220^+^ cells than Lpr mice, suggesting reduced antibody-producing plasma lineage cells and reduced GC B cell activities (supplementary Figure 2D). mTORC2 is known to promote Tfh cell differentiation (18). We observed a reduced CXCR5^+^PD-1^hi^ cell population in Lpr-*Ifnar1^−/−^* mice (Figure 3G). Lpr-*Ifnar1^−/−^* mice did not show significantly reduced ICOS expression on CD4^+^ T cells (supplementary Figure 2E) but had a significantly lower percentage of CXCR5^+^Bcl6^+^ and CXCR5^+^Bcl6^-^ populations than Lpr mice (supplementary Figure 2G), reminiscent of the phenotypes of Lpr mice with T cell specific deletion of *Rictor* (16). mTORC2 activation is also known to suppress Treg function (19). However, Treg frequency was not significantly altered in Lpr-*Ifnar1^−/−^*mice compared to Lpr mice (supplementary Figure 2F), suggesting that type I IFN signaling might not substantially affect Treg homeostasis in Lpr mice. Together, our results demonstrated that IFNAR1 deficiency partially rescues autoimmune phenotypes in Lpr mice, associated with reduced systemic inflammation, Tfh differentiation and B cell activation, associated with reduced mTORC2 activity. Our data reaffirms the disease promoting function of type I IFN in Lpr mice and provides evidence of association between type I IFN and mTORC2 signaling in SLE.

**Figure 3:**
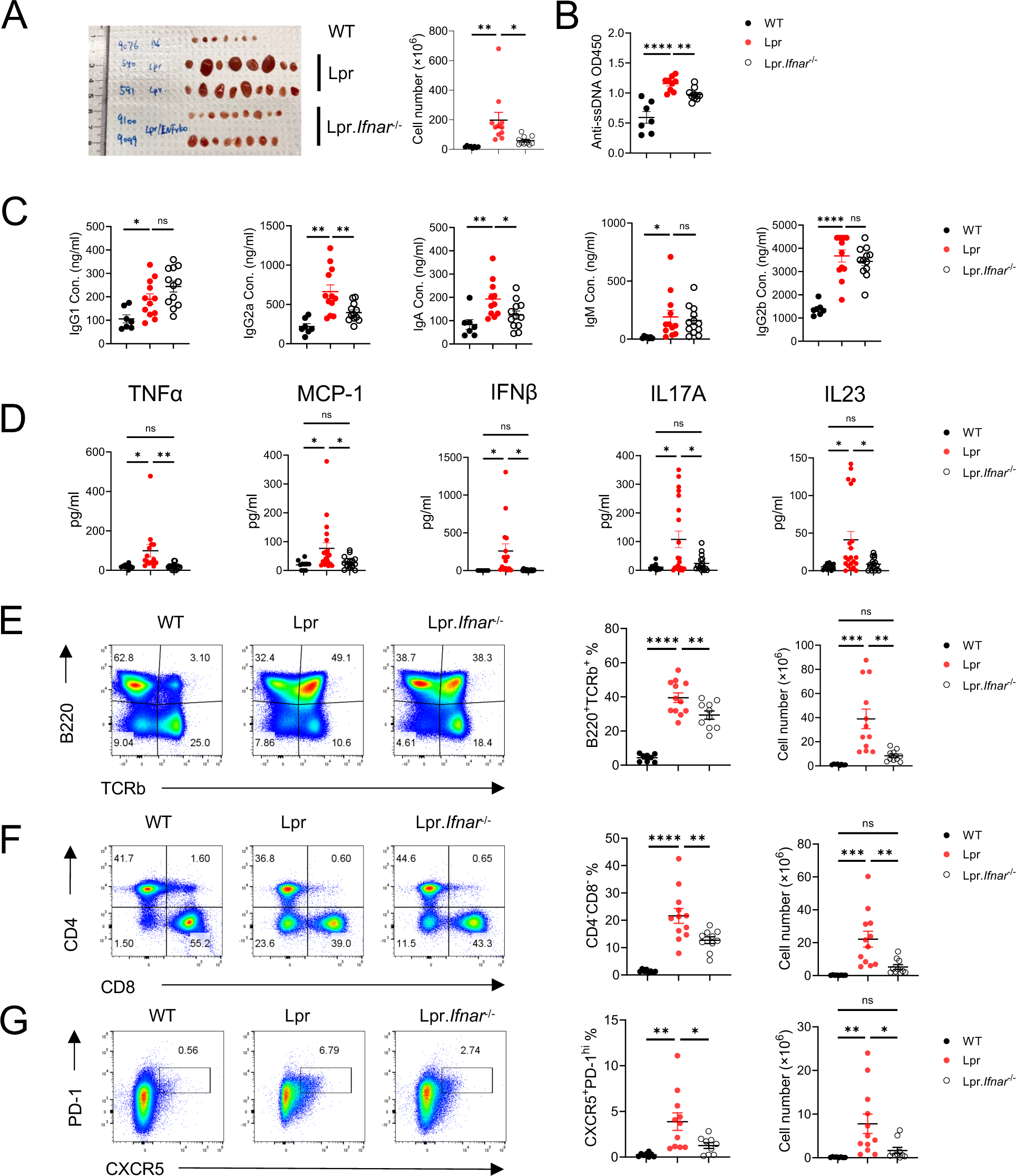
IFNAR1 deletion reduces systemic inflammation in Lpr mice. (A) Representative image of peripheral lymph nodes (left) and cellularity (right) from 6 months B6, Lpr, and Lpr*.Ifna1r^-/-^* mice. (B) Tiers of anti-ssDNA antibody level in mice serum measured by ELISA. Serum immunoglobulin (Ig) isotype concentration (C) and inflammatory cytokine concentration (D) in 6-month-old mice. (E) Expression of TCRb and B220 in pLN derived lymphocytes. Left, frequency of TCRb^+^B220^+^ population; right absolute cell number of TCRb^+^B220^+^ cells in pLN. (F) Expression of CD4 and CD8 in pLN derived lymphocytes. Left, frequency of CD4^-^CD8^-^ population; right absolute cell number of CD4^-^CD8^-^ cells in pLN. (G) Expression of CXCR5 and PD-1 in pLN derived lymphocytes. Left, frequency of CXCR5^+^PD-1^hi^ Tfh cells; right, absolute cell number of CXCR5^+^PD-1^hi^ Tfh cells in pLN. ns, not significant; *p < 0.05, **p < 0.01, ***p < 0.001, ****p < 0.0001. p-Values were calculated with one-way ANOVA with post hoc Tukey test. Error bars represent SEM.

### *Rictor*-ASO treatment benefits lupus-like symptoms in MRL/*lpr* mice

After demonstrating that genetic targeting mTORC2 in T cells ameliorates SLE in mouse models and the close link between mTORC2 and type IFN signaling in SLE development, we sought to target mTORC2 pharmacologically. We designed antisense oligonucleotides (ASOs) that specifically targets mouse *Rictor*. Immunoblot analysis showed that anti-mouse *Rictor*-ASO could effectively delete RICTOR and abrogate AKT phosphorylation at S473 site in mouse CD4^+^ T cells without affecting mTORC1 signaling (Figure 4A). We next tested if *Rictor*-ASO exhibited beneficial effects in MRL/*lpr* mice (Figure 4B), a mouse strain with spontaneous lupus-like symptoms commonly used for SLE therapeutic testing. We observed that *Rictor*-ASO could significantly improve the survival probability of MRL/*lpr* mice (Figure 4C) and reduce proteinuria levels in those mice (Figure 4D) than Ctrl-ASO. At 19 weeks, *Rictor*-ASO treated mice had lower creatinine concentration in both urine (Figure 4E) and serum (Figure 4F), as well as reduced serum urea nitrogen concentration (Figure 4G) than in Ctrl-ASO treated mice, suggesting improved renal function. *Rictor*-ASO treated mice also had smaller pLN (supplementary Figure 3A). Flow analysis showed that *Rictor*-ASO treatment increased the percentage of CD4^+^ T cells in pLN of MRL/*lpr* mice (supplementary Figure 3B). Decreased CD69 expressing CD4^+^ T cells were also observed in *Rictor*-ASO treated mice (supplementary Figure 3C), consistent with our data from genetic models. We also detected lower autoantibody (Figure 4H) and immunoglobulin isotype levels in serum (Figure 4I) in *Rictor*-ASO treated mice, indicating reduced humoral immune activation. Reduced levels of inflammatory cytokines in *Rictor*-ASO treated mice suggested reduced systemic inflammation (Figure 4J). Finally, kidney histology revealed that *Rictor*-ASO treated mice have a lower number of foci and a lower level of cortex inflammation than Ctrl-ASO treated mice at 19 weeks (Figure 4K), indicating less extent of nephritis. Taken together, these data indicate that *Rictor*-ASO treatment can enhance survival and ameliorate lupus-like diseases in MRL/*lpr* mice.

**Figure 4:**
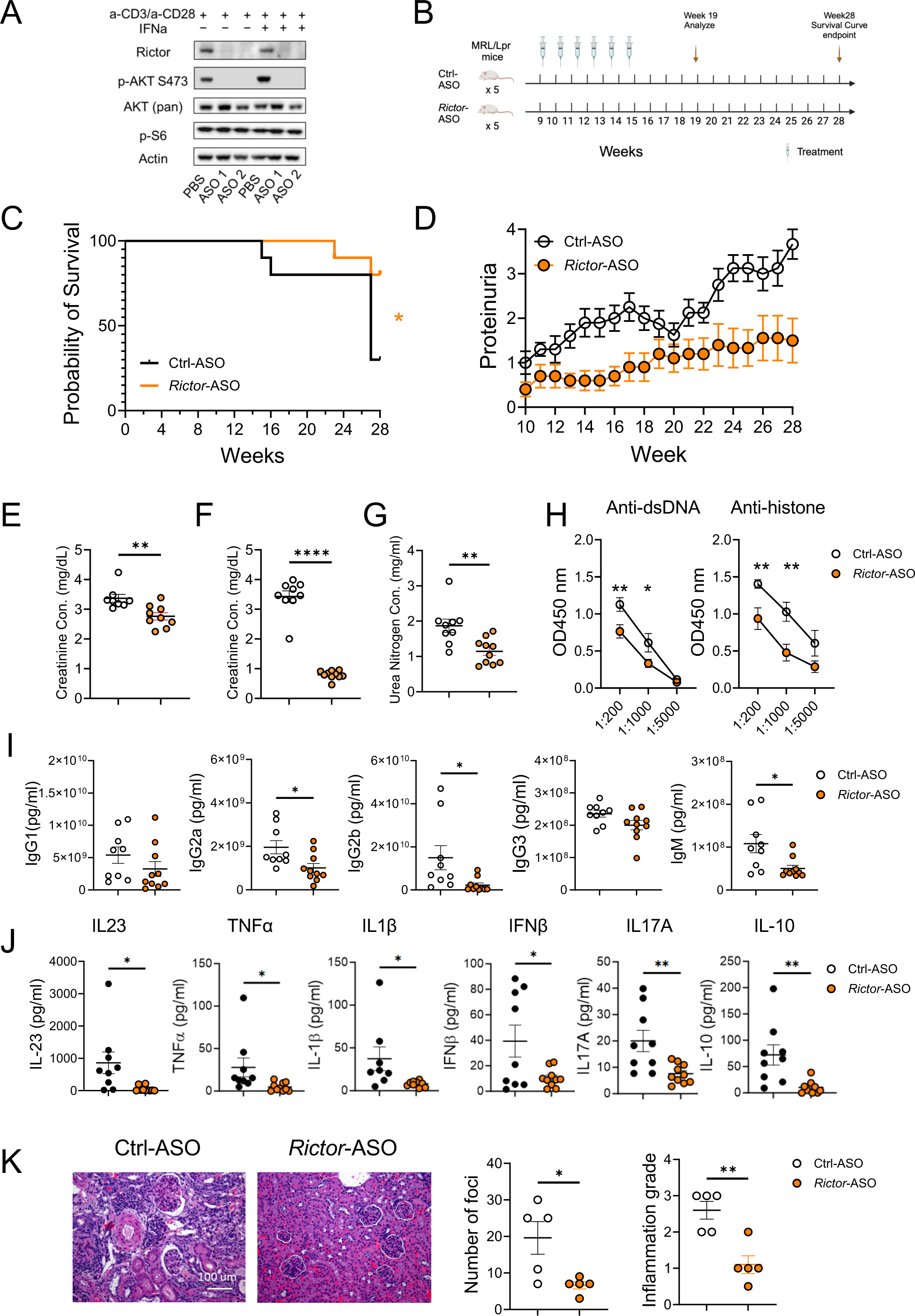
Anti-mouse *Rictor*-ASO treatment benefits lupus-like symptoms in MRL/*lpr* mice. (A) Immunoblot image showing the deletion of RICTOR by *Rictor*-ASO in mouse CD4^+^ T cells. (B) Treatment scheme of *Rictor-*/Ctrl-ASO in MRL/*lpr* mice. (C) Kaplan–Meier cumulative survival plot of *Rictor*-ASO and Ctrl-ASO treated MRL/*lpr* mice. (D) Proteinuria changes of *Rictor*-ASO and Ctrl-ASO treated MRL/*lpr* mice over 28 weeks. Urine (E) and serum (F) creatinine concentration of MRL/*lpr* mice at 19 weeks. (G) Serum urea nitrogen concentration at 19 weeks. (H) Tiers of anti-dsDNA (left) and anti-histone (Right) antibodies in mouse serum at 19 weeks. Serum inflammatory cytokine concentration (I) and Ig isotypes concentration (J) at 19 weeks. (K) Representative H&E staining images of kidneys at 19 weeks; Left: quantitative cortex inflammation grade; right: number of foci. Scale bar: 100 μm. ns, not significant; *p < 0.05, **p < 0.01. The Kaplan–Meier estimator was used for survival curve analysis (C). Unpaired student t-tests were used for two groups comparison (E – K). Error bars represent SEM.

### *RICTOR*-ASO treated SLE patient derived Tfh-like cells reduce antibody production in T:B coculture assay

Aberrant mTORC2 activation and increased circulating Tfh-like cells have been shown to strongly correlate with disease activity in SLE patients (32,33). We finally asked if targeting mTORC2 using human *RICTOR*-ASO in human SLE Tfh cells can reduce B cell activities *in vitro*. We showed that *RICTOR*-ASO could effectively reduce RICTOR expression and mTORC2 activity in human CD4^+^ T cells (Figure 5A). To test whether inhibition of mTORC2 in Tfh cells can reduce Tfh mediated antibody production *in vitro,* we sorted Tfh-like cells (CD4^+^CXCR5^+^PD-1^+^CXCR3^-^) from SLE patients’ (53.6 ± 4.15 years) PBMCs. After treating sorted Tfh cells with *RICTOR*-/Ctrl-ASO for 5 days, memory B cells (CD19^+^IgD^-^CD27^+^ CD38^-^) sorted from the same donor-matched SLE PBMCs were cocultured with ASO-treated Tfh cells for 7 days (Figure 5B). SLE patients’ demographics and disease characteristics were shown in Supplementary Table 1. At day 7, we observed substantially lower immunoglobulin isotypes (i.e. IgG1, IgG3, IgG4, IgM) in the supernatant of *RICTOR*-ASO treated SLE T-B cocultures than those of the Ctrl-ASO counterparts (Figure 5C). Reduced IgG1 and IgG4, but not other, immunoglobulin level was also seen in *RICTOR*-ASO treated T-B coculture derived from age-matched healthy donor (HC, 55 ± 5.01 years) PBMCs (supplementary Figure 4). Autoantigen array showed significantly reduced autoantibodies signal against various autoantigens, such as Lo/SSO and Ra/SSA, in *RICTOR*-ASO treated SLE T-B coculture compared to paired Ctrl-ASO counterparts (Figure 5D, E). These data suggest that targeting mTORC2 in SLE patient Tfh cells could effectively reduce immunoglobulin isotypes and autoantibodies production, highlighting a promising therapeutic potential in clinics.

**Figure 5:**
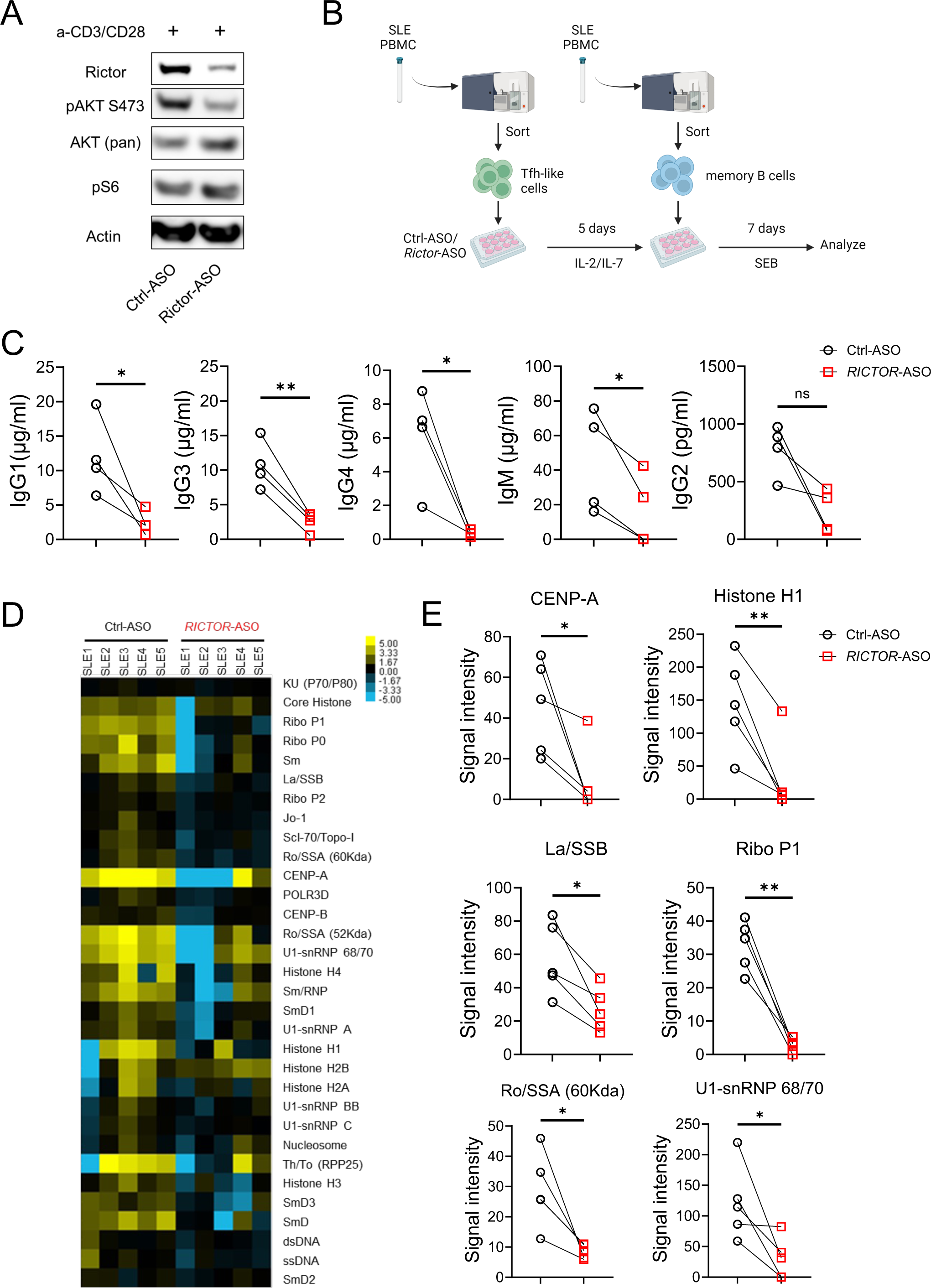
Anti-human *RICTOR*-ASO reduces antibody production in an *in vitro* T-B cell coculture assay. (A) Immunoblot showing the deletion of RICTOR in human CD4^+^ T cells. (B) Experiment scheme of the *in vitro* T-B cell coculture assay. (C) Ig isotypes concentration in coculture supernatant after 7 days. (D) Signal intensity heatmap of antibodies against autoantigens in culture supernatant. (E) Paired analysis of autoantibodies signals detected in culture supernatant. ns, not significant; *p < 0.05, **p < 0.01. p-Values were calculated with paired t-tests. Error bars represent SEM.

## Discussion

In this study, we tackled the contribution of mTORC2 in T cells for lupus disease development, investigated the relationship between mTORC2 and type I IFN in SLE pathogenesis, and explored the therapeutic potential of mTORC2 targeting ASOs in mice and humans. We showed that genetic deletion of RICTOR in mice can effectively ameliorate IMQ-induced lupus-like diseases. We also found that the loss of IFNAR1 in Lpr mice correlates with reduced mTORC2 activities. Finally, we developed novel anti-mouse *Rictor-* and anti-human *RICTOR-*ASO that efficiently and specifically suppress mTORC2. Anti-mouse *Rictor*-ASO can delay disease onset and benefit lupus nephritis in MRL/*lpr* mice. Inhibiting RICTOR expression in human SLE patient derived Tfh-like cells by anti-human *RICTOR*-ASO can reduce immunoglobulin isotypes and autoantibodies production in an *in vitro* T-B coculture system. These results showed that mTORC2 plays a pivotal role in SLE disease development and interconnected with known molecular pathways in SLE pathogenesis. Therefore, targeting the mTORC2 pathway using ASOs could be an effective therapeutic for future SLE management.

Both type I and type II IFN signaling dysregulation have been reported in human SLE (34). However, how IFN signaling regulates CD4^+^ T cell differentiation and trafficking in SLE is not fully understood. Gain-of-function genetic variants in both type I and II IFN pathways have been identified as risk factors for SLE (35,36). Type I IFN signaling has been recognized as a key player in SLE pathogenesis with elevated IFNα and IFNg gene signatures as hallmarks of human SLE (37), and anti-IFN-α/β receptor antibodies can reduce lupus-like symptoms in mice (14) and humans (38). One of the intriguing findings from a large-scale single cell RNAseq analysis of 162 SLE patients is that type I IFN signature is highly correlated with CD4^+^ T cell lymphopenia, correlates with the known function of type I IFN-CD69 axis on T cell egress (30,39). Consistent with this observation, treatment with anifrolumab, an IFNAR1 antagonist, is associated with correction of T cell lymphopenia symptom in SLE patients (40). Our investigation builds on earlier study and provides potential immunological mechanisms through which type I IFN may contribute to SLE. Our data corroborates with the clinical observation that type I IFN activation partially contributes to CD4^+^ T cell lymphopenia, overactivation of Tfh, GC and extrafollicular responses. Importantly, type I IFN activation is also associated with mTORC2 activation and increased T cell glucose metabolism in Lpr mouse model. Indeed, the immunological phenotypes of Lpr-*Ifnar1^−/−^*mice closely resemble those of Lpr-*Rictor* T-KO mice, including the partial rescue of CD69 expression, CD4^+^ T cell lymphopenia, Tfh/GC differentiation (without affecting ICOS expression on T cells) and T cell glucose metabolism. Overall, the magnitude of rescue is stronger in Lpr-*Rictor* T-KO mice than in Lpr-*Ifnar1^−/−^* mice, suggesting that factors other than IFNAR1 signal through mTORC2 in Lpr mice. These results highlighted the close relation between mTORC2 signaling and classic SLE pathogenic signaling, further indicating its indispensable role in SLE development.

While mTORC2 activities are known to elevate in human SLE (41), the lack of selective mTORC2 inhibitors has long hindered functional studies and therapeutic development. RNA-based silencing technique showed the feasibility of specific silencing of RICTOR and corresponding therapeutic effects in a breast cancer model (42), pointing the direction of mTORC2-based therapies. ASOs have emerged as the new generation of therapies for a wide range of rare diseases such as neurological, inherited metabolic, and infectious diseases (43). ASO targeting mTORC2 was first successfully used to ameliorate a neurological disorder induced by brain-specific loss of PTEN (44). Our results showed that mTORC2 targeting ASOs can effectively and specifically suppress mTORC2 in both human and mouse CD4^+^ T cells. Administration of *RICTOR*-ASO in MRL/*lpr* mice can improve mice survival, kidney function, and immunopathology, suggesting its promising therapeutic potential. Further, we showed that reducing RICTOR expression by anti-human *RICTOR*-ASO in human SLE Tfh cells can reduce immunoglobulin and autoantibodies production when cocultured with donor-matched memory B cells. These results further support the idea of using *RICTOR*-ASOs for human SLE management. Given the heterogeneity of human SLE, it is crucial to map the mTORC2 signature in larger-scale patient populations in future studies to enable individualized mTORC2-based therapy. Given the relatively narrow immunological impact of mTORC2 inhibition compared to mTORC1 inhibition (19), it is plausible that targeting mTORC2 specifically could have fewer immune suppressive side effects than mTORC1 inhibition. Future comparative studies are warranted to address this question.

In summary, our study provides mechanistic insight that associates mTORC2 with type I IFN signaling in lupus immunopathology. We further provide proof-of-principle evidence that ASO mediated mTORC2 inhibition could ameliorate lupus symptoms using lupus mouse model and *in vitro* T-B cell co-culture derived from SLE patients. Future pre-clinical and clinical studies will be needed to identify mTORC2 signatures in heterogeneous SLE patients and test the safety and efficacy of ASO treatment for human SLE.

## Acknowledgment

We acknowledge Mayo Clinic Arizona Histology Core Laboratory for the process and histological staining of mouse kidney specimens. We acknowledge Genecopoeia Inc. for autoantigen array analysis. The study is supported by NIH R01AR077518 and R01AI162678 (to H.Z.), lupus Research Alliance (696599, to H.Z.) and Mayo Foundation for Medical Education and Research.

## Author contributions

All the authors were involved in drafting or revising the article critically for important intellectual content, and all authors approved the final version to be published. Dr. Zeng has full access to all the data in the study and takes responsibility for the integrity of the data and the accuracy of the data analysis. M.A. and H.Z. conceived the study, designed the experiments and wrote the manuscript. M.A., X.Z. and Y.L. prepared the research material and carried out the experiments. M.C. and P.J. designed and provided the control, anti-*Rictor* and anti-*RICTOR* ASOs. M.A.B. and A.A.D.G analyzed patient clinical data. N.G. and M.A. performed the pathological review.

## Supplementary Methods

### Flow cytometry

The following antibodies were used in flow cytometry: anti-B220 (RA3-6B2), anti-CD4 (GK1.5), anti-CD8a (53-6.7), anti-CD25 (PC16), anti-CD38 (90), anti-CD69 (H1.2F3), anti-GL7 (GL-7), anti-CD138 (281-2), anti-IgD (11-26c.2a), anti-CD95 (Jo2), anti-PD-1 (J43), anti-IgM (II/41), and anti-CD162 (2PH1). CXCR5 and PNA were stained with biotinylated anti-CXCR5 (2G8) or biotinylated peanut agglutinin (FL10-71), followed by staining with streptavidin-conjugated PE (BD Biosciences).

### SCIENTH method

Cells were rested in complete medium at 37°C for 30 minutes before equally divided into four parts and seeded into 96 well plates. Wells were treated with vehicle or the following metabolic inhibitors for 15 minutes, 2-Deoxy-D-Glucose (2-DG, 100 mM), Oligomycin (Oligo,1 mM), or a sequential combination of the two. Puromycin (10 µg/ml) was then added to each treated well for 15 minutes. Cells were then washed with ice-cold PBS and stained with Fc receptors and viability dye at RM for 15 minutes. Cells were then stained with surface markers in FACS buffer at RM for 20 minutes. Following washing, cells were fixed and permeabilized using the FOXP3 fixation and permeabilization kit (Biolegend) following the manufacturer’s instructions. Cells were next stained with anti-Puromycin AF647 (Sigma Aldrich, clone 12D10), resuspended in FACS buffer and read on an Attune NxT (ThermoFisher) cytometer.

### T-B coculture assay

CD4+ T cells were first enriched from the healthy or SLE donor PMBC using an enrichment kit (STEMCELL, Cat# 17952). The B220-CD3+CD4+CXCR5+PD-1+CXCR3-cells were sorted as Tfh-like cells. Tfh cells (3×104 cells/well) were cultured with anti-human RICTOR- or Ctrl-ASO (10 nm), IL2 (100 U/ml), and IL7 (10 ng/ml) for 5 days. CD19+ B cells were next enriched from same donor PBMC using an enrichment kit (STEMCELL, Cat# 17854). CD19+IgD-CD27+ CD38-cells were next sorted as memory B cells (2×104 cells/well) and seeded to coculture with ASO treated Tfh cells in the presence of staphylococcal enterotoxin B (SEB, 100ng/ml, Toxin Technology) for 7 days.

**Supplementary Table 1.** Demographic and clinical characteristics of SLE patients*. *All patients met ACR/EULAR SLE criteria. Abbreviations: ANA=antinuclear antibodies; dsDNA=anti-double-stranded DNA antibody; RNP=anti-Ribonucleoprotein antibody. Sm=anti-Smith antibody; Ro=anti-Ro antibody; La=anti-La antibody; SCL70= anti-topoisomerase I; RF= Rheumatoid Factor. ACCP= Anti-cyclic citrullinated peptide, aCL = anticardiolipin; anti-β2GPI = anti–β2-glycoprotein I, HCQ= Hydroxychloroquine.

| **Patient** | **1** | **2** | **3** | **4** | **5** |
| --- | --- | --- | --- | --- | --- |
| **Age (years)** | 57 | 36 | 55 | 67 | 53 |
| **Gender** | Female | Male | Female | Female | Female |
| **Race/Ethnicity** | White non-Hispanic | White non-Hispanic | White non-Hispanic | White non-Hispanic | White non-Hispanic |
| **SLE duration (Years)** | 28 | 12 | 7 | 12 | 19 |
| **SLEDAI 2k Score** | 9 | 10 | 10 | 4 | 0 |
| **Autoantibodies** |  |  |  |  |  |
| **ANA** | + | + | + | + | + |
| **Anti-Sm** | - | + | - | + | - |
| **Anti-RNP** | + | - | - | + | - |
| **Anti-Ro** | + | - | - | - | - |
| **Anti-La** | + | - | - | - | - |
| **Anti-dsDNA** | + | + | - | - | - |
| **aCL IgM / IgG** | - | Not available | Not available | +/+ | - |
| **β2GP1 I IgM / IgG** | - | Not available | Not available | - | - |
| **Lupus anticoagulant** | - | Not available | Not available | - | + |
| **Complement C3/4** | Low | Normal | Normal | Normal | Normal |
| **Organ Involvement** |  |  |  |  |  |
| **Joints** | + | + | + | + | + |
| **Constitutional** | - | - | - | - | - |
| **Hematologic** | + | - | - | - | + |
| **Mucocutaneous** | - | + | + | - | + |
| **Kidney** | - | - | - | - | - |
| **Serosal** | + | - | - | - | - |
| **Neuropsychiatric** | - | - | - | - | - |
| **Treatments** |  |  |  |  |  |
| **Glucocorticoids** | + | + | - | + | - |
| **HCQ** | + | - | + | + | + |
| **Mycophenolate** | - | - | - | - | - |
| **Methotrexate** | - | - | + | + | - |
| **Azathioprine** | - | - | - | + | + |
| **Others** | None | None | None | None | None |

\*All patients met ACR/EULAR SLE criteria.
Abbreviations: ANA=antinuclear antibodies; dsDNA=anti-double-stranded DNA antibody; RNP=anti-Ribonucleoprotein antibody. Sm=anti-Smith antibody; Ro=anti-Ro antibody; La=anti-La antibody; SCL70= anti-topoisomerase I; RF= Rheumatoid Factor. ACCP= Anti-cyclic citrullinated peptide, aCL = anticardiolipin; anti-β2GPI = anti-β2-glycoprotein I, HCQ= Hydroxychloroquine.

### Supplementary figure legend

**Supplementary Figure 1:**
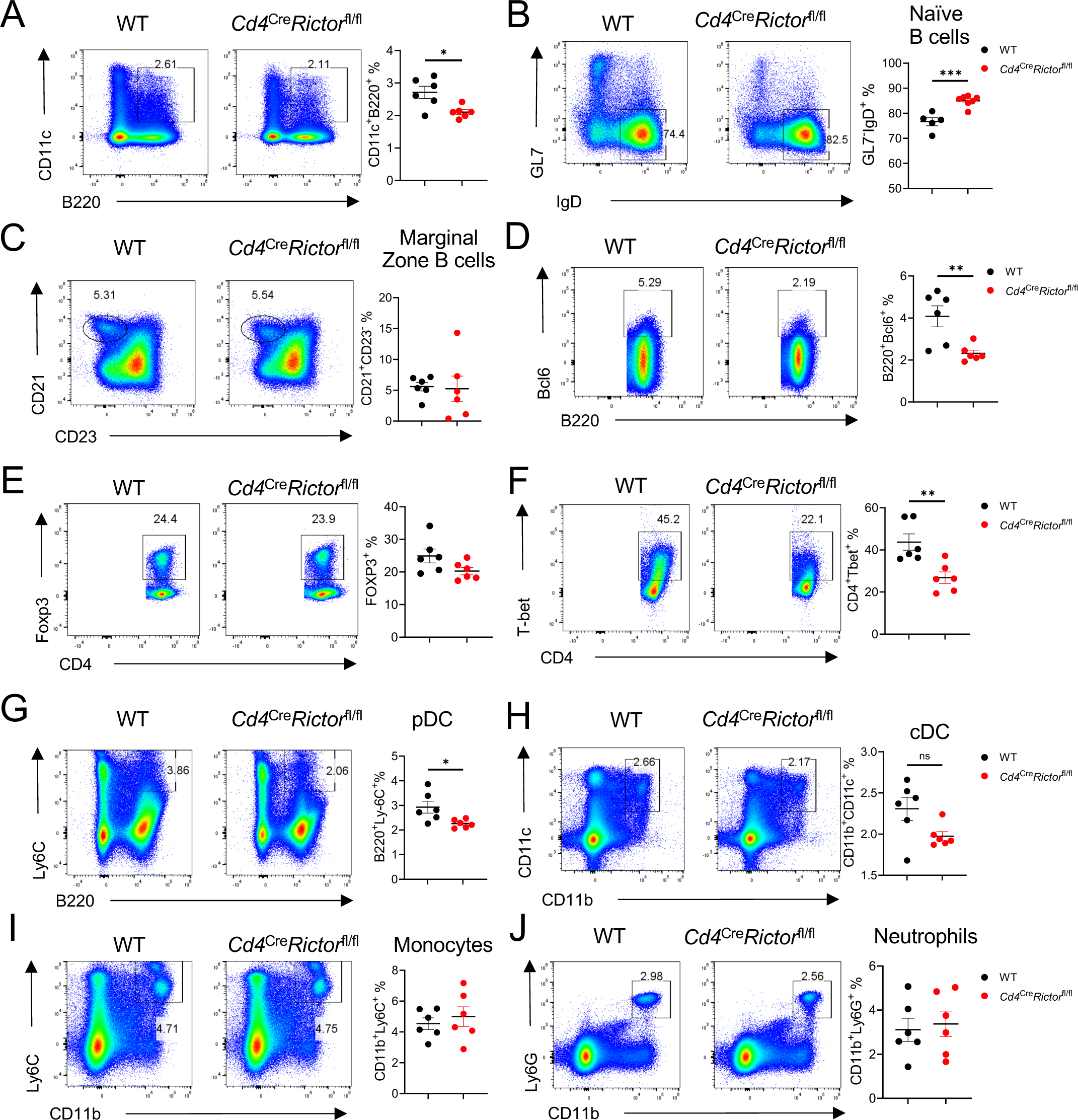
Flow analysis of splenocyte cell populations in IMQ mice. (A) Expression of GL7 and IgD in splenocyte. Right, frequency of GL7^-^IgD^+^ naïve B cells. (B) Expression of CD21 and CD23 in splenocyte. Right, frequency of CD21^+^CD23^-^ marginal zone B cells. (C) Expression of Bcl6 and B220 in splenocyte. Right, frequency of B220^+^Bcl6^+^ cells. (D) Expression of CD11c and B220 in splenocyte. Right, frequency of CD11c^+^B220^+^ cells. (E) Expression of Foxp3 and CD4 in splenocyte. Right, frequency of Foxp3^+^ cells within CD4^+^ cells. (F) Expression of T-bet and CD4 in splenocyte. Right, frequency of CD4^+^T-bet^+^ cells within CD4^+^ cells. (G) Expression of Ly6C and B220 in splenocyte. Right, frequency of B220^+^Ly6C^+^ pDC cells. (H) Expression of CD11c and CD11b in splenocyte. Right, frequency of CD11c^+^ CD11b^+^ cDC cells. (I) Expression of Ly6C and CD11b in splenocyte. Right, frequency of CD11b^+^Ly6C^+^ monocytes. (J) Expression of Ly6G and CD11b in splenocyte. Right, frequency of CD11b^+^Ly6G^+^ neutrophils. *p < 0.05, **p < 0.01, ***p < 0.001. p-Values were calculated with unpaired student t-tests. Error bars represent SEM.

**Supplementary Figure 2:**
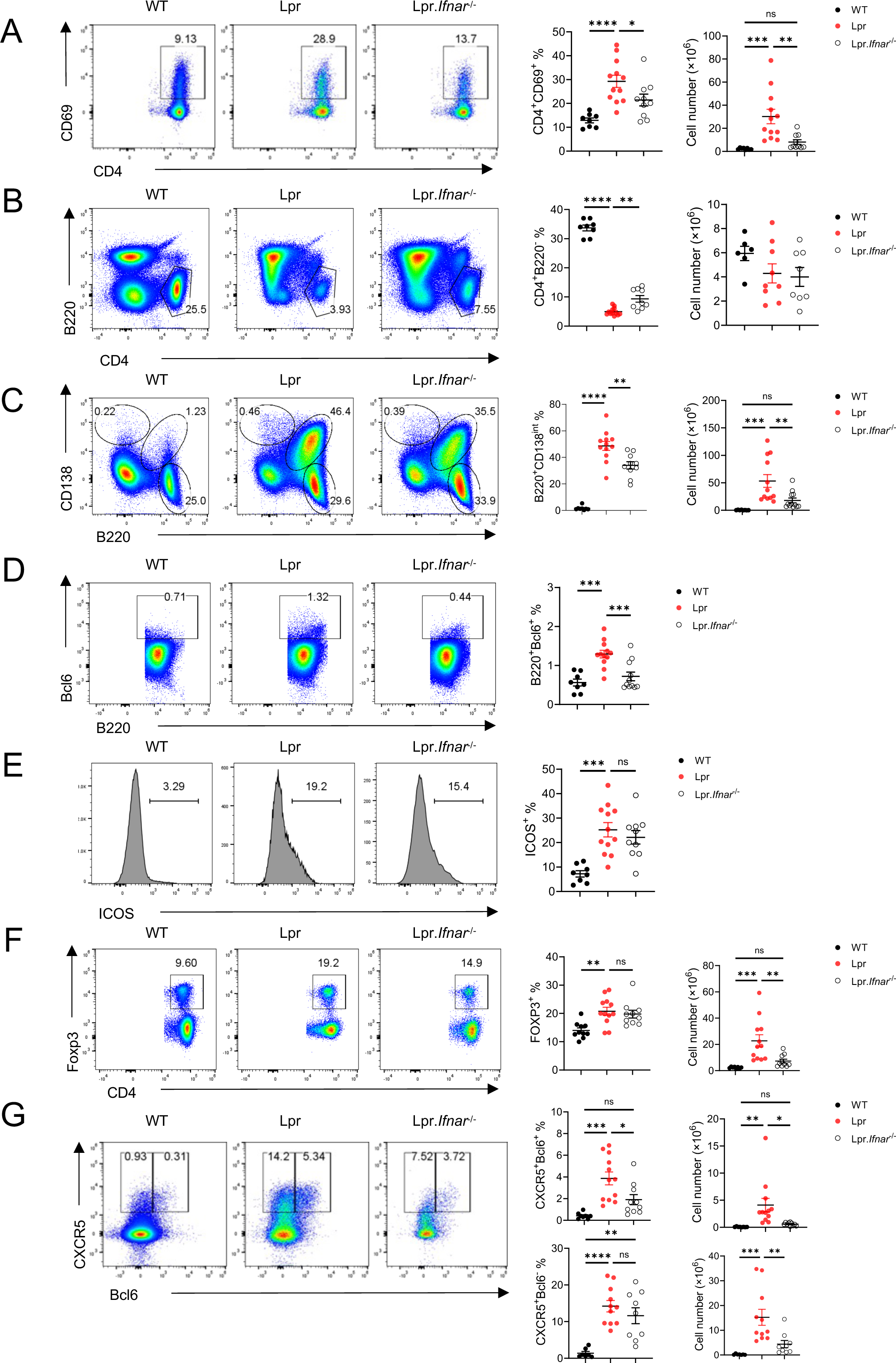
Flow analysis of peripheral lymphocytes in Lpr-*Ifnar1^-/-^* mice. (A) Expression of CD69 and CD4 in pLN derived lymphocytes. Left, frequency of CD4^+^CD69^+^ population; right absolute cell number of CD4^+^CD69^+^ cells in pLN. (B) Expression of B220 and CD4 in pLN derived lymphocytes. Left, frequency of CD4^+^B220^-^ population; right absolute cell number of CD4^+^B220^-^ cells in pLN. (C) Expression of B220 and CD138 in pLN derived lymphocytes. Left, frequency of B220^+^CD138^hi^ population; right absolute cell number of B220^+^CD138^hi^ cells in pLN. (D) Expression of B220 and Bcl6 in pLN derived lymphocytes. Right, frequency of B220^+^Bcl6^+^ population in pLN. (E) Expression of ICOS on CD4^+^ cells in pLN. (F) Expression of Foxp3 and CD4 in pLN derived lymphocytes. Left, frequency of CD4^+^Foxp3^+^ population; right absolute cell number of CD4^+^Foxp3^+^ cells in pLN. (G) Expression of CXCR5 and Bcl6 in pLN derived lymphocytes. Upper left, frequency of CXCR5^+^Bcl6^+^ population; upper right, absolute cell number of CXCR5^+^Bcl6^+^ cells in pLN. Lower left, frequency of CXCR5^+^Bcl6^-^ population; lower right, absolute cell number of CXCR5^+^Bcl6^-^ cells in pLN. Results were pooled from at least 3 independent experiments. ns, not significant; *p < 0.05, **p < 0.01, ***p < 0.001, ****p < 0.0001. p-Values were calculated with one-way ANOVA with the post-hoc Tukey test. Error bars represent SEM.

**Supplementary Figure 3:**
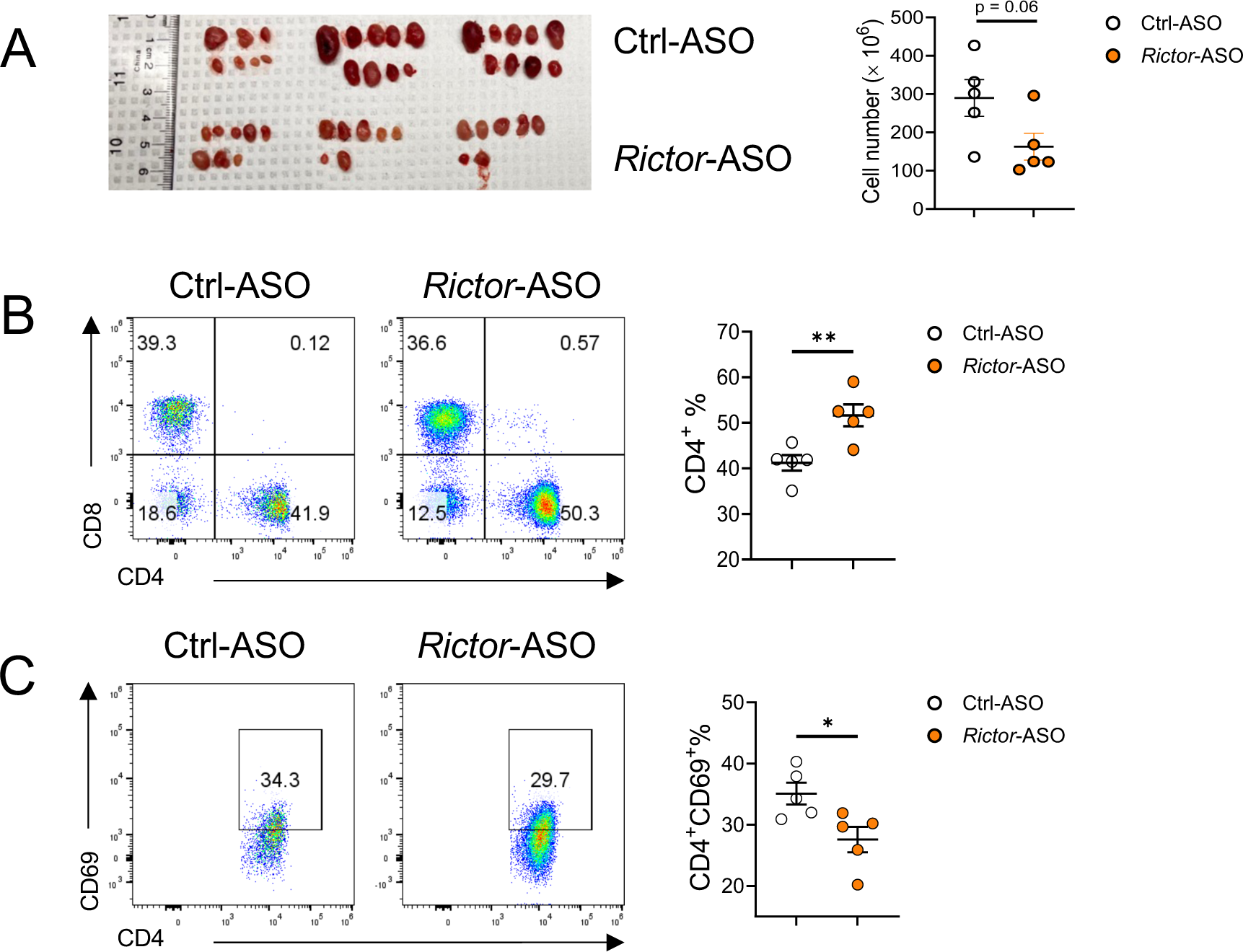
Flow analysis of peripheral lymphocytes in Ctrl-/*RICTOR*-ASO treated MRL/*lpr* mice. (A) Peripheral lymph node size (left) and derived lymphocyte cell numbers (Right) in treated mice. (B) Expression of CD4 and CD8 in pLN derived lymphocytes. Right, percentage of CD4^+^ population in pLN. (C) Expression of CD4 and CD69 in pLN derived lymphocytes. Right, frequency of CD4^+^CD69^+^ population in pLN. *p < 0.05, **p < 0.01. p-Values were calculated with unpaired student t-tests. Error bars represent SEM.

**Supplementary Figure 4:**
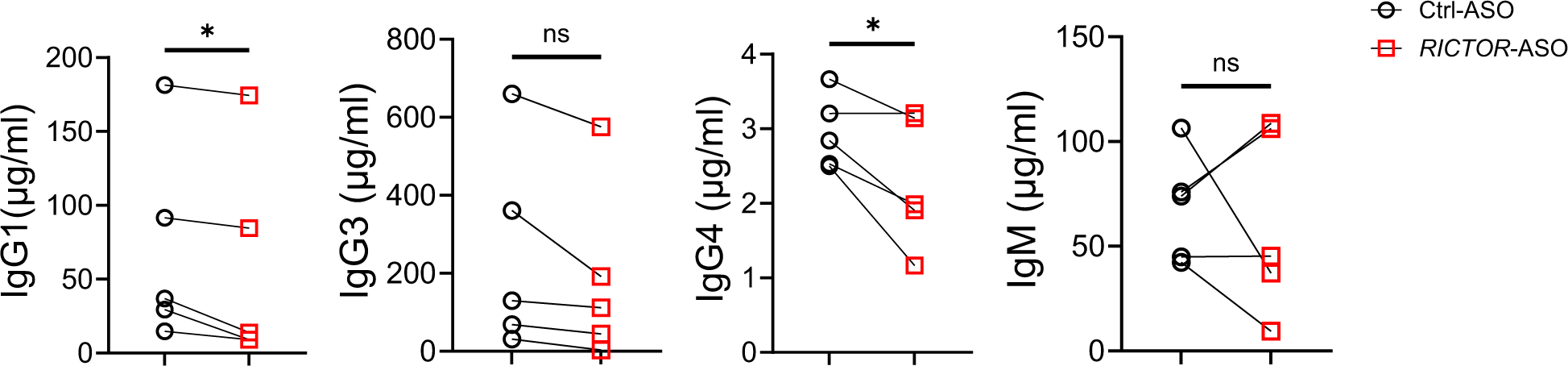
Supernatant immunoglobulin isotypes concentration of T-B culture derived from healthy donor PBMCs. ns, not significant, *p < 0.05, **p < 0.01. p-Values were calculated with paired t-tests. Error bars represent SEM.

